# A screen of alkaline and oxidative formulations for their inactivation efficacy of metal surface adsorbed prions using a steel-bead seed amplification assay

**DOI:** 10.1101/2023.06.26.546570

**Authors:** Daniel Heinzer, Merve Avar, Manuela Pfammatter, Rita Moos, Petra Schwarz, Matthias T. Buhmann, Benjamin Kuhn, Stefan Mauerhofer, Urs Rosenberg, Adriano Aguzzi, Simone Hornemann

## Abstract

Iatrogenic transmission of prions, the infectious agents of fatal Creutzfeldt-Jakob disease, through inefficiently decontaminated medical instruments remains a critical issue. Harsh chemical treatments are effective, but not suited for routine reprocessing of reusable surgical instruments in medical cleaning and disinfection processes due to material incompatibilities. The identification of mild detergents with activity against prions is therefore of high interest but laborious due to the low throughput of traditional assays measuring prion infectivity. Here, we report the development of TESSA (s**T**ainl**ES**s steel-bead **S**eed **A**mplification assay), a prion seed amplification assay that explores the propagation activity of prions with stainless steel beads. TESSA was applied for the screening of about 70 different commercially available and novel formulations and conditions for their prion inactivation efficacy. One hypochlorite-based formulation, two commercially available alkaline formulations and a manual alkaline pre-cleaner were found to be highly effective in inactivating prions under conditions simulating automated washer-disinfector cleaning processes. The efficacy of these formulations was confirmed *in vivo* in a murine prion infectivity bioassay, yielding a reduction of the prion titer for the bead surface adsorbed prions below detectability. Our data suggest that TESSA represents an effective method for a rapid screening of prion-inactivating detergents, and that alkaline and oxidative formulations are promising in reducing the risk of potential iatrogenic prion transmission through insufficiently decontaminated instrument surfaces.

## Introduction

Transmissible spongiform encephalopathies (TSEs), or prion diseases, represent a group of fatal disorders which are hallmarked by the misfolding and aggregation of the cellular prion protein (PrP^C^) into a pathological, proteinase K (PK) resistant form, PrP^Sc^ (Prusiner 1982). Prions can occur as different strains that are characterized by different inheritable physicochemical properties and clinical signs of the diseases (Caughey, Raymond et al. 1998). It is widely accepted that the different strain properties are encoded in the different conformations of the prion agent (Aguzzi, Heikenwalder et al. 2007). Prion diseases can be inherited, or arise sporadically as Creutzfeldt-Jakob disease (sCJD) in humans (Aguzzi and Calella 2009). In addition, the nature of the infectious agent, the prion, allows manifestation of the disease through exposure to infected tissue via ingestion or through iatrogenic transmission (Aguzzi and Calella 2009).

The first recorded case of iatrogenic CJD (iCJD) as a result of a corneal transplant operation dates back to 1974 (Duffy, Wolf et al. 1974). Later reports have shown that prion diseases are transmissible through contact with contaminated surgical devices, dura-mater transplants, and human growth hormone administration (Bernoulli, Siegfried et al. 1977, Collins and Masters 1996, Bonda, Manjila et al. 2016). The decontamination of prions poses a major challenge, as they are incredibly resistant to common cleaning, disinfection and sterilization processes (Lehmann, Pastore et al. 2009, Hughson, Race et al. 2016, Nakano, Akamatsu et al. 2016) which were originally developed for the inactivation of viruses and microorganisms (Alper, Cramp et al. 1967, Zobeley, Flechsig et al. 1999, Taylor 2000, McDonnell and Burke 2003). Moreover, prions strongly adhere to a broad variety of surface materials (Zobeley, Flechsig et al. 1999, Lipscomb, Pinchin et al. 2006, Luhr, Low et al. 2009, Secker, Hervé et al. 2011, Mori, Atarashi et al. 2016), which further complicates their removal and decontamination by conventional cleaning and instrumental reprocessing routines, leaving a potential risk of transmission of prion diseases through insufficiently decontaminated medical devices. The World Health Organization (WHO) has therefore published a recommendation list of strong chemical and physical procedures that should be applied for the decontamination of prions (WHO 2000). Such procedures, however, are not applicable to routine, daily medical device reprocessing or to non-disposable medical devices, such as endoscopes and surgical instruments, due to material incompatibilities (Head and Ironside 2007). Also, preventive measures such as the utilization of single-use tools, or allocating instruments only for the use on potential CJD cases, are not reasonable solutions, as they would lead to insurmountable costs with questionable feasibility (Thomas, Chenoweth et al. 2013). Hence, there is still a need for mild prion decontamination products that can be applied for the reprocessing of surgical instruments.

To date, the efficacy of novel prion decontaminants is routinely assessed in rodent bioassays that report on prion infectivity based on the survival of indicator mice inoculated with prions (Flechsig, Hegyi et al. 2001, Peretz, Supattapone et al. 2006, Giles, Glidden et al. 2008, Berberidou, Xanthopoulos et al. 2013). However, rodent assays are limited in throughput, cost-intensive and take several months to complete. In addition, *in vivo* bioassays should be replaced by *in vitro* assays, where possible, in compliance with the 3R of animal welfare. The recent development of *in vitro* seed amplification assays (SAA), such as the protein misfolding cyclic amplification (PMCA) (Saborio, Permanne et al. 2001) and real-time quaking induced cyclic amplification assay (RT-QuIC) (Wilham, Orrú et al. 2010, Frontzek, Pfammatter et al. 2016) that take advantage of the seeding capabilities of prions allow to conduct several experiments in parallel in a much shorter time with higher throughput.

In this study, we report the establishment of a SAA, termed TESSA (s**T**ainl**ES**s steel-bead **SA**A), that uses stainless steel beads as prion carriers, to mimic the steel surface of surgical instruments. We applied TESSA for a screen of a series of formulations and conditions to explore their prion-inactivating (prionicidal) capacity for automated medical device reprocessing. A hypochlorite-based formulation was identified as the most efficient prion decontaminant in the screen and was further validated together with two commercially available alkaline formulations and an alkaline pre-cleaner in a mouse bioassay. All products reduced prion titers below detectability in the bioassay, demonstrating the usefulness of these formulations as potential prion decontaminants and TESSA as a valuable tool for the rapid evaluation of novel prionicidals.

## Results

### Establishment of TESSA

To develop an assay that allows for the fast screening of novel anti-prion decontaminants, we modified a microplate SAA (Atarashi, Sano et al. 2011, Frontzek, Pfammatter et al. 2016) for the detection of surface-attached prions using micron-sized stainless-steel beads as prion carriers (AISI 316L). The prion-exposed steel beads were first tested for their capability to efficiently bind prions. Beads were exposed to either 1% (w/v) prion-infected mouse brain homogenate (RML6, Rocky Mountain Laboratory strain, passage 6; 9.9 ± 0.2 Log LD50 units g^-1^; (Falsig, Julius et al. 2008)) or non-infectious brain homogenate (NBH), followed by air-drying to strengthen the adherence of the prions to the metal beads. After intense washing with ddH2O, eluates of RML6 and NBH coated beads were analyzed by immunoblotting for surface bound PrP^C^ and proteinase K (PK)-resistant PrP^Sc^, as a marker for the presence of prions (McKinley, Bolton et al. 1983). Total PrP was detectable in eluates of non-digested RML6 and NBH coated beads (Fig. 1A), confirming efficient binding of PrP to the steel-beads. After PK digestion, PrP^C^ was completely digested in the NBH reference sample and on NBH coated beads, whereas RML6 displayed the typical electrophoretic mobility pattern of the PK-resistant core fragments of PrP^Sc^ (Bolton, Meyer et al. 1985, McKinley, Meyer et al. 1991). Eluates from PK-digested RML6 coated beads, however, showed a decreased signal intensity and no changes in the electrophoretic distribution pattern when compared to the undigested beads. This finding suggests that the PK cleavage sites of PrP^Sc^ may have become protected due to the adsorption of PrP^Sc^ to the beads, so that PrP^Sc^ largely remained preserved as full-length protein on the beads.

**Figure 1:**
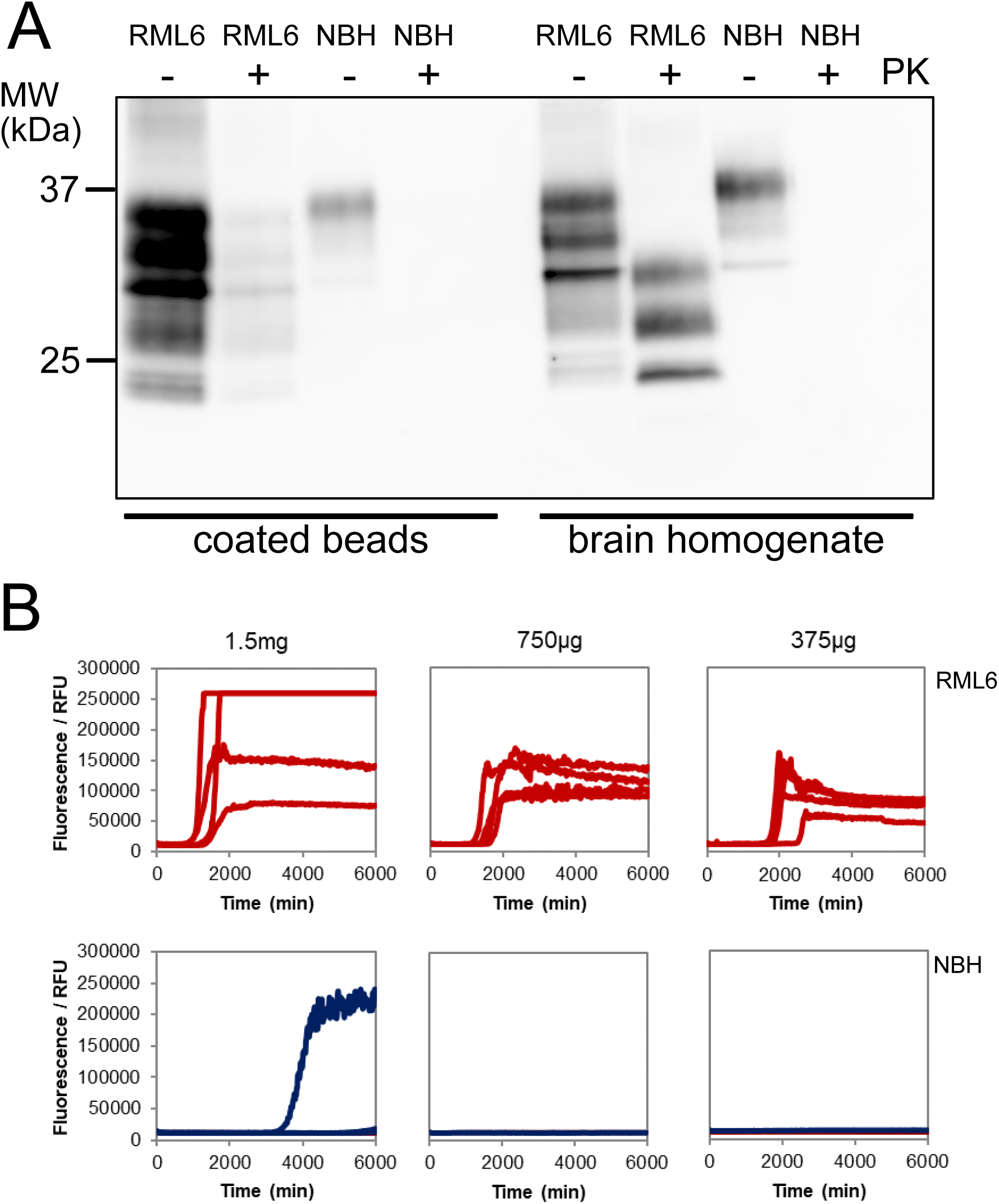
Evaluation of PrP^Sc^-specific binding to the stainless-steel beads. **(A)** PK-immunoblot analysis demonstrating the binding of total PrP and PrP^Sc^ to the steel beads. Eluates from NBH and RML6 exposed beads were blotted after treating the samples without (-) or with (+) PK for the detection of total PrP and PrP^Sc^. As controls RML6 and NBH without and with PK digestion were loaded. The anti-PrP antibody POM-1 was used for detection. The molecular weight standard is shown in kilodaltons (kDa). **(B)** TESSA showing specific propagation activity for three different bead amounts of RML6 coated beads as indicated in the Figure. NBH coated beads were used as negative controls. All reactions were performed in quadruplicates.

To further assess whether prions, upon adhering to the beads, retained their characteristic propagation activities and to identify the maximum amount of beads with the highest specific sensitivity in the TESSA, we analyzed the seeding properties of three different amounts of beads coated with either RML6 or NBH in quadruplicate reactions (1.5 mg, 750 µg and 375 µg/TESSA reaction, Fig. 1B). Efficient seeding was detected for all amplification reactions of RML6-coated beads (Fig. 1B). These data demonstrate that prions retained their propagation activity after adsorption to the beads. For the NBH-coated beads, one non-specific positive reaction out of four replicates was detected at the highest amount of 1.5 mg/TESSA reaction. To avoid any potential risk of non-specific positive reactions and technical difficulties in the handling of the highest bead amount due to the high viscosity of the bead suspension, we chose 750 µg beads per TESSA reaction as the optimal amount for the formulation screening.

### Applicability of TESSA as a method for testing prion decontaminants

To investigate the applicability of TESSA for the assessment of new prion decontaminants, we first evaluated its performance with 1 M NaOH (2 h; RT), a common standard decontamination method for the inactivation of prions (WHO 2000), and three commercially available alkaline formulations. These formulations included (Table 1): (i) deconex^®^ 28 ALKA ONE-x (28AO); an alkaline cleaner whose alkalinity is solely based on potassium metasilicate and that has been shown to be effective for the decontamination of the Chandler strain of prion diseases *in vitro* (Hirata, Ito et al. 2010), (ii) 28AO in combination with deconex^®^ TWIN ZYME (28AO/TZ); a mild alkaline product with improved cleansing properties through the enzymatic activities of subtilisin and α-amylase, and (iii) deconex^®^ 36 BS ALKA (36BS); a non-enzymatic highly alkaline pre-cleaner that was presumed to exhibit its prion decontamination efficacy due to a pH-value of >11, though still being applicable to a broad range of materials despite its relatively high pH-value.

**Table 1.** Summary of the chemical composition of the lead formulations and conditions used to decontaminate the prion-contaminated steel beads.

| Formulation | Ingrediens | Experimental conditions |
| --- | --- | --- |
| 36BS | - Tripotassium orthophosphate (15-30 %)<br>- Anionic surfactants (< 5 %)<br>- Non-ionic surfactants (< 5 %)<br>- Alanine, N,N-bis(carboxymethyl) sodium salt (5-15 %) | Concentration: 2 % in water with standardized hardness<br>Condition: 25 °C, 60 min |
| 28AO | - Tripotassium orthophosphate (15-30 %)<br>- Dipotassium Trioxosilicate (5-15 %)<br>- Amphoteric surfactants (< 5 %) | Concentration: 1 % in ddH<sub>2</sub>O<br>Condition: 70 °C, 10 min |
| 28AO/TZ | - N,N-Dimethyldecylamine N-oxide (1-5 %)<br>- Subtilisin (< 1 %)<br>- Alpha-Amylase (< 1%)<br>- Non-ionic surfactants (15-30 %)<br>- Potassium sorbate (5 %) | Concentration: 1 % 28AO + 0.3 % TZ in ddH<sub>2</sub>O<br>Two step reprocessing<br>1. 45 °C, 10 min<br>2. 70 °C, 10 min |
| 70B | - Tetrapotassium pyrophosphate (15-30 %)<br>- Dipotassium trioxosilicate (5-15 %)<br>- Sodium hypochlorite solution, active chlorine (< 5 %) | Concentration: 1 % in ddH<sub>2</sub>O<br>Condition: 55 °C, 10 min |

Beads were exposed to RML6 (1 % (w/v)), intensively washed, and treated with either NaOH or the three different formulations 28AO, 28AO/TZ or BS under conditions recommended by the manufacturer (Table 1). After another intense washing step to remove any residual decontaminant, beads were analyzed by TESSA. NaOH and all three formulations efficiently inactivated the seeding activity of the prion exposed beads below detectability (Fig. 2A and, Fig. S1A), whereas beads treated with deionized water (ddH2O) maintained their prion seeding characteristics. This data indicate that TESSA is a suitable *in vitro* tool for the screening and effectiveness testing of novel prion decontaminants and that formulations 28AO, 28AO/TZ, and 36BS have promising anti-prion inactivation activities.

**Figure 2:**
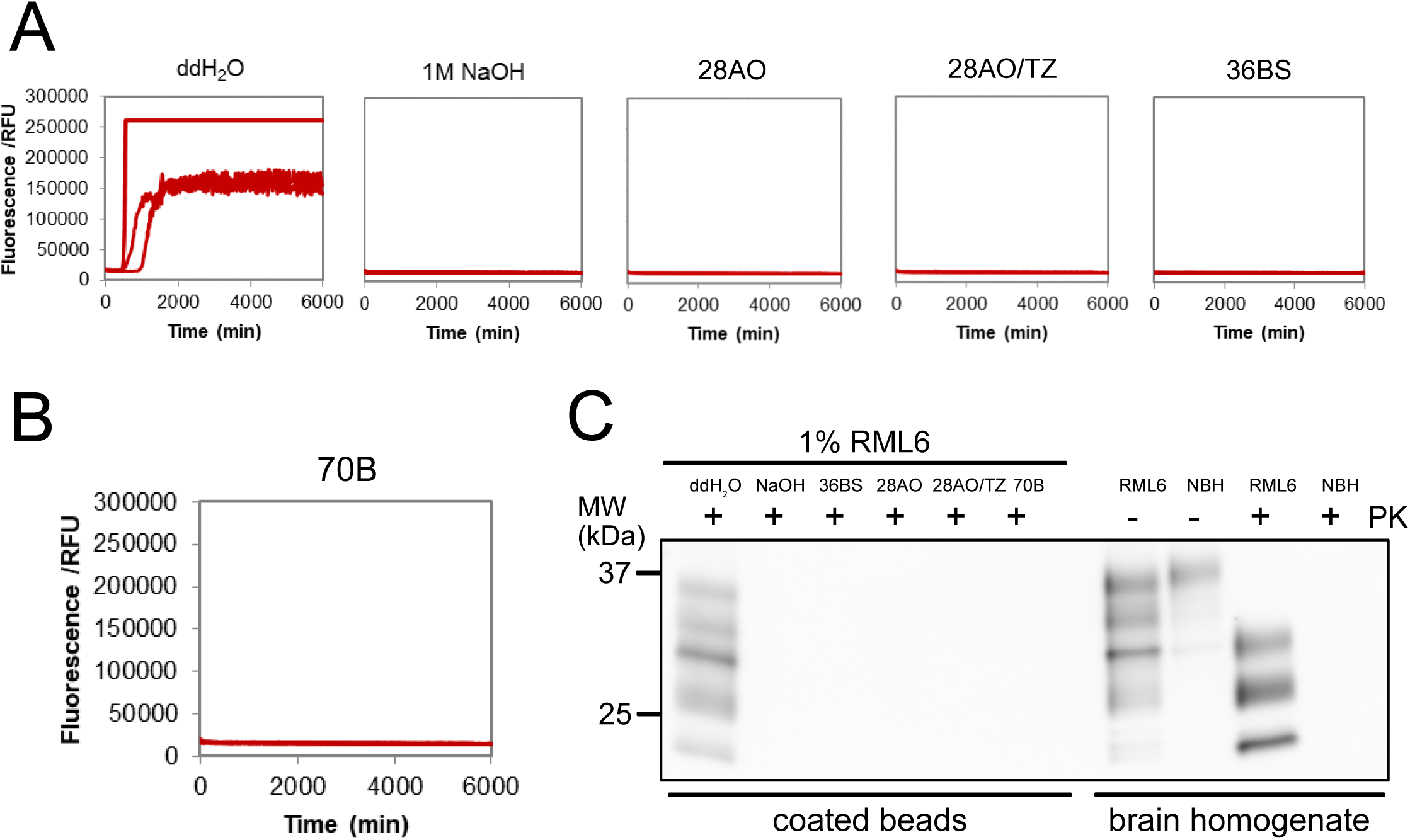
TESSA and PK-immunoblot analysis to assess the decontamination effectiveness of the four lead formulations. **(A)** TESSA analysis of RML6 exposed and treated beads. No remaining seeding activity was observed for RML6 exposed beads after decontamination with the different formulations. RML6 exposed beads treated with ddH2O and 1 M NaOH were used as positive and negative controls, respectively. Shown is one representative data set in quadruplicates for each condition. **(B)** Same as (A), but after decontamination with 70B. **(C)** PK-immunoblot showing no residual PrP^Sc^ in the eluates of RML6 exposed beads after treatment with the four formulations. As further controls, RML6 and NBH before (-) and after PK (+) digestion are shown.

### Screening of different formulations for their prion inactivation effectiveness using TESSA

We then applied TESSA to screen about 70 different mildly alkaline and oxidative formulations and conditions for their anti-prion efficacy (summarized in Table 2). Based on their main active ingredient, the different formulations were grouped into 4 different categories post-hoc. The first category comprised formulations, for which the anti-prion effect was expected to be mediated by the hydrolytic activity of alkanolamines and complexing agents, which are common donors of alkalinity in detergent solutions with a typical pKa of around 9.5. The second category included alkaline enzymatic phosphate/silicate-based formulations (TWIN PH10). The alkalinity of these products is based on the phosphate/silicate composition with the advantage of higher material compatibility. Category 3 encompassed a selection of commercially available strong alkaline hydroxide-and hydroxide/silicate-based formulations and solutions (Category 3, Table 1), whereas formulations of category 4 were composed of hypochlorite (70B); a chemical that acts as a strong oxidizing reagent under alkaline conditions (Sandin, Karlsson et al. 2015). To support the anti-prion activity and cleaning properties of the main components, in particular, of products of Categories 1 and 2, these were formulated with a series of enzymatic, non-ionic surfactants and surface-active reagents (Table 2).

**Table 2:**
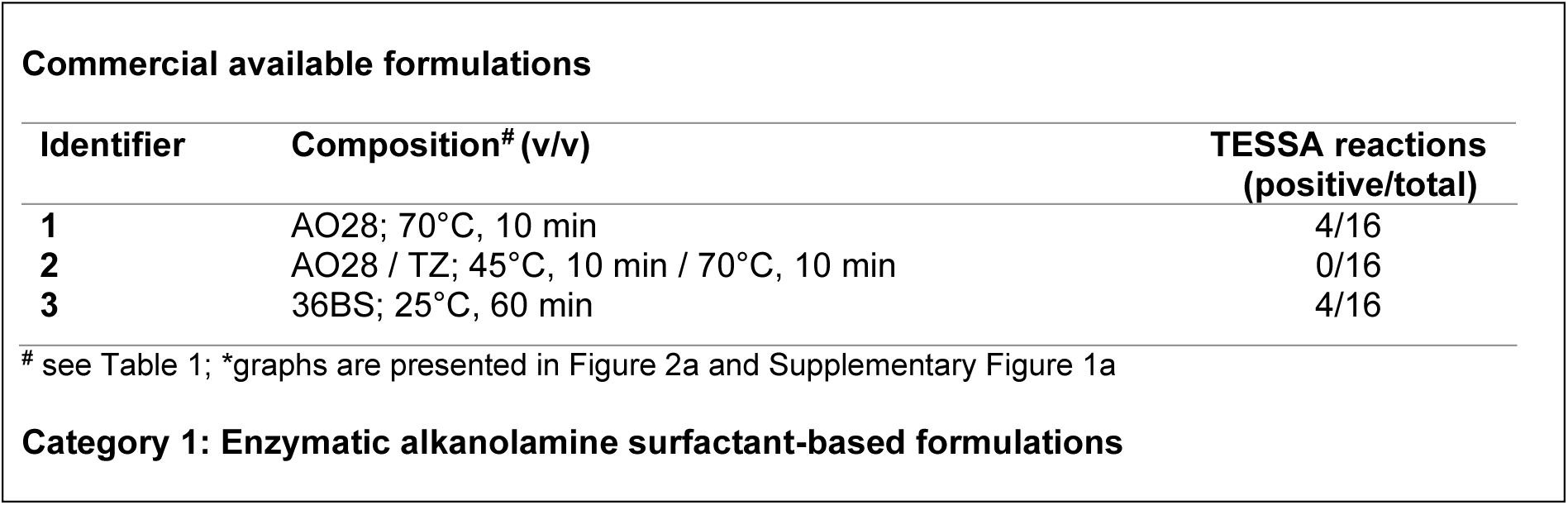
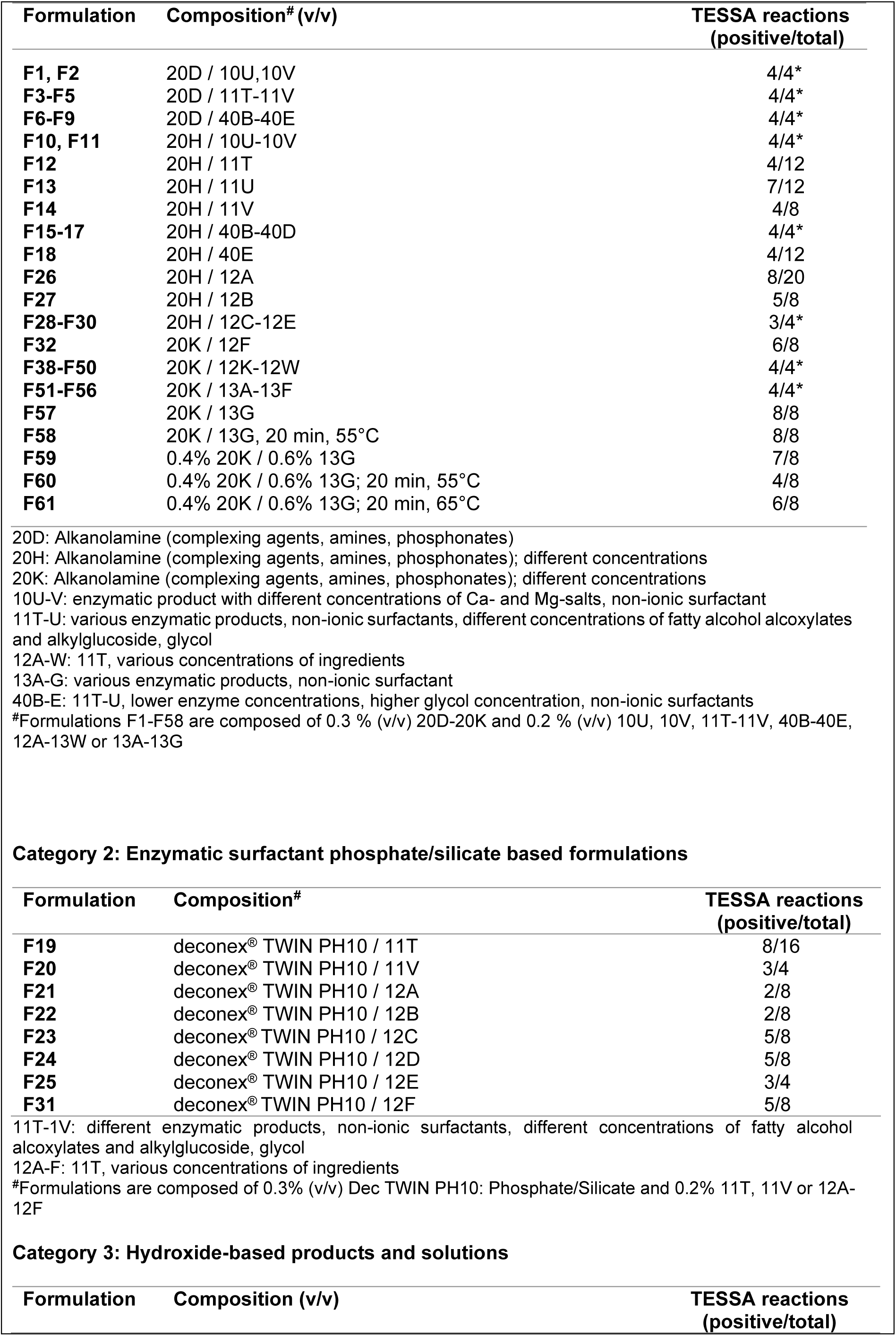
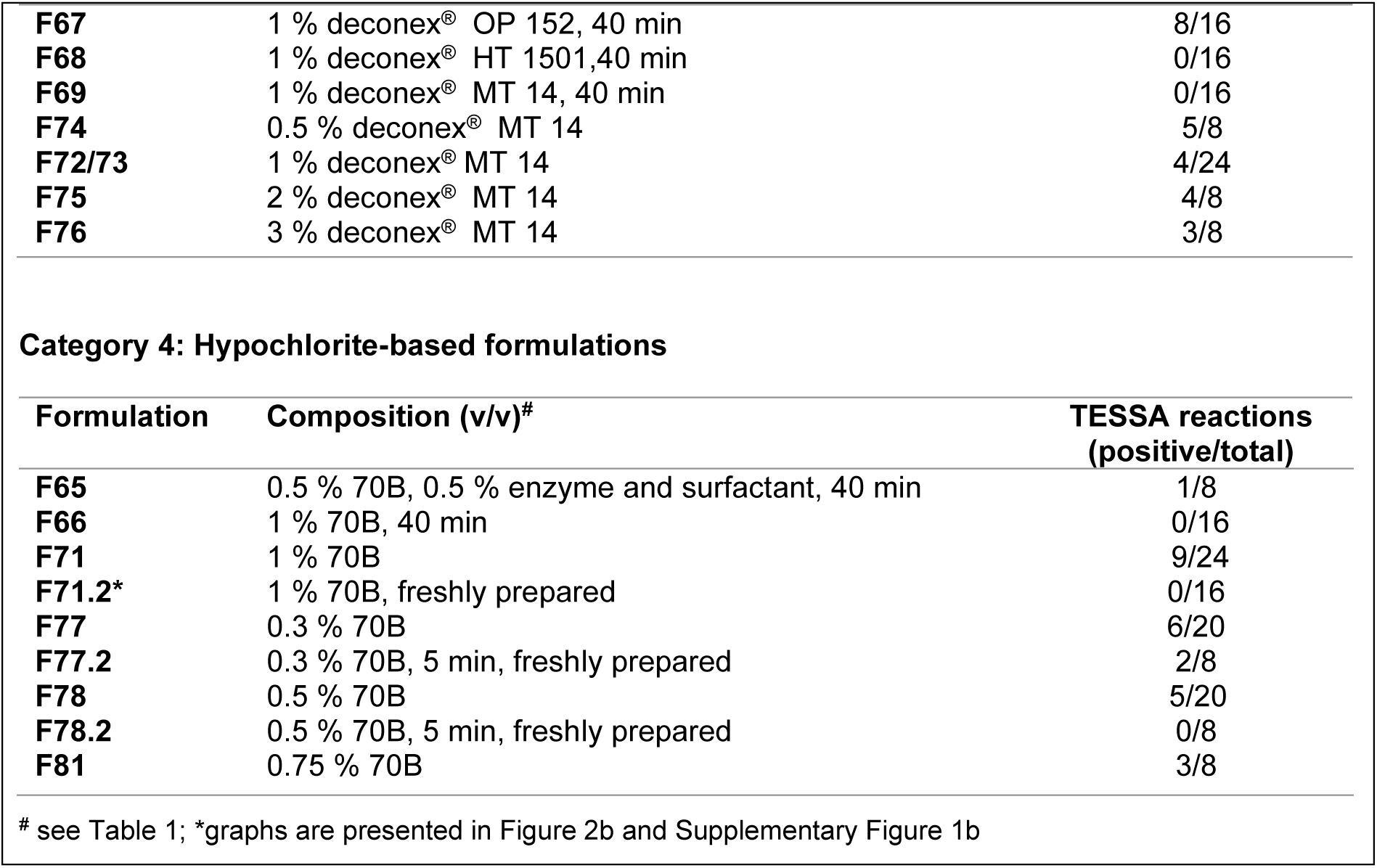
Post-hoc classification of the formulations and experimental conditions. The number of positive replicates from up to 5 independent TESSA experiments with 4 replicates each are shown. All formulations were applied under machine reprocessing conditions (55 °C, 10 min), if not otherwise indicated. *Number of positive/total replicate for each formulation.

The effectiveness of the different formulations was studied under commonly used automated reprocessing conditions as used in washer-disinfectors (10 min, 55 °C). Some formulations were also applied for longer treatment times (40 min and 2 h) and at further temperatures to assess their capability to inactivate prion seeding activity under differing conditions. Beads were treated as described above and analyzed by TESSA with the number of replicates as indicated in Table 2. Although some formulations reduced the prion seeding activity below detectability at extended treatment times (40 min and 2 h), most formulations showed only moderate to no inactivation efficiency when applied at 55 °C for 10 min. The most effective formulation under this condition was the hypochlorite-based formulation 70B. Although the data for 70B showed some variability over time, which we attributed to the degradation of hypochlorite due to the limited shelf life of chlorine containing products (Nicoletti, Siqueira et al. 2009), when prepared freshly, 70B was able to remove all detectable prion seeding activity (Fig. 2B, Fig. S1B and Table 1). Formulation 70B was also effective at an even lower concentration and shorter contact time of 0.5 % (v/v) and 5 min (Table 2). We therefore conclude that formulation 70B exhibits the most efficient prion decontamination capability among the tested products, but with the limitation of the stability of hypochlorite containing formulations (Clarkson, Moule et al. 2001, Nicoletti, Siqueira et al. 2009, Hughson, Race et al. 2016).

### Efficacy confirmation by immunoblotting and in a mouse bioassay

We then selected formulations 70B, 28AO, 28AO/TZ, and 36BS to further assess their prion decontamination effectiveness. We first analyzed the eluates of prion-coated beads after treatment with the different formulations and NaOH for the presence of PrP^Sc^ by PK immunoblotting (Fig. 2C). After treatment, no residual PrP^Sc^ was detectable on the immunoblot, showing that all decontamination products effectively reduced the prion load on the stainless-steel beads.

To investigate if the identified formulations were also able to reduce prion infectivity on the prion-coated beads, we further confirmed their decontamination capability in *tg*a*20* mice, a transgenic mouse line that overexpresses murine PrP^C^ (Fischer, Rulicke et al. 1996). We first determined the sensitivity and maximal titer reduction that can be achieved in the bioassay after decontamination of the prion-coated beads by performing an end-point dilution titration. Three independent batches of 25 mg beads were incubated in 100 µL of 10-fold serial dilutions of RML6 (10^-2^ – 10^-7^; Table 3). After intensive washing with ddH2O and air-drying, beads were resuspended in 1 mL PBS. Bead suspensions of 30 µL (750 µg beads/mouse) were then inoculated intracerebrally (i.c.) into three groups of three *tg*a*20* mice each (n= 9 per dilution, from three independent bead preparations). The survival of the inoculated mice was monitored for 250 days post infection (dpi). Only mice inoculated with beads exposed to RML6 dilutions from 10^-2^ to 10^-4^ developed clinical signs of a prion disease with complete attack rates at 75.9 ± 1.8, 92.8 ± 5.5 and 121 ± 13.2 dpi, respectively (Fig. 3A, Table 3). To further confirm that mice succumbed to prion disease, brains of the mice (10^-2^ dilution of RML6) were extracted and immunohistologically investigated for the presence of spongiform changes and for the accumulation of PrP^Sc^ (Fig. 3B). Data were analyzed by comparison to a standard curve obtained from an end-point titration of RML6 inoculated *tg*a*20* mice (Falsig, Julius et al. 2008) and a median 50 % lethal dose [LD50] of about 5.1 log10 infectious units ml^-1^ was estimated for the prion-exposed bead suspension inoculated per mouse (Table 3). This value represents the maximal sensitivity achievable in the mouse bioassay and is similar to the prion titer that has previously been determined in mice for individual steel wires (10^5.5^ LD50 wire units) coated with either RML (Edgeworth, Sicilia et al. 2011) or the 263K hamster strain (Lemmer, Mielke et al. 2008, McDonnell, Dehen et al. 2013).

**Figure 3:**
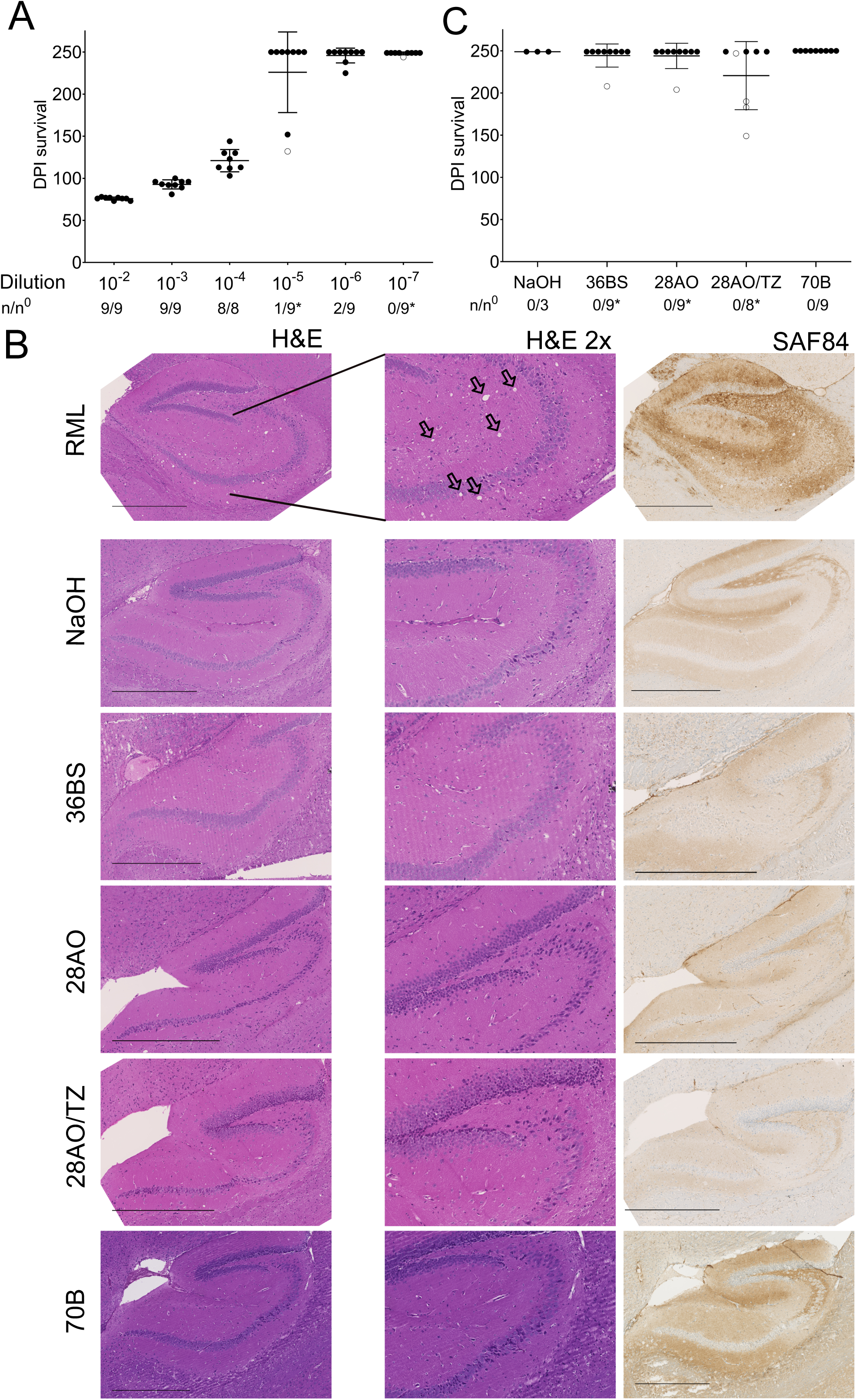
Mouse bioassay for the evaluation of the decontamination efficiency of the different formulations. **(A)** End-point titration in *tg*a*20* mice inoculated with beads exposed to 10-fold serial dilutions of RML6 from 10^-2^ to 10^-7^. Data points are shown as the mean incubation time ± standard deviation of the mean. n/n^0^: indicates the attack rate (number of mice developing a prion disease divided by the total number of inoculated mice). Open circles: individual mice that died of an intercurrent death unrelated to prion infection. **(B)** Histopathology of brain sections from *tg*a*20* mice inoculated with RML6 exposed steel beads treated with either ddH2O, 1 M NaOH or the different formulations (as indicated in the Figure). Vacuolation and PrP^Sc^ deposits as markers for the presence of prion disease were observed after visualization by H&E staining (middle: expansion, 2 × magnification) and imaging with an antibody directed against PrP (SAF84), whereas no vacuolation and PrP^Sc^ deposits were observed in the brains mice inoculated with the treated beads. Vacuoles are indicated by black arrows. (Scale bars: 500 µm). **(C)** Survival of *tg*a*20* mice inoculated with RML6 exposed beads treated either with 1M NaOH or the different formulations. None of the mice developed a prion disease until the end of the experiment.

**Table 3.** Serial dilutions of RML6 coated beads to estimate the titer of the RML6 coated beads in an end-point format and efficacy testing of the different formulations on the RML6 coated beads in a mouse bioassay (Falsig, Julius et al. 2008). All formulations reduced the prion infectivity titer by at least 5.1 log10. Treatment with 1 M NaOH and the different formulations was performed on beads coated with a 10^-2^ dilution of RML6. ^a^ Data for the end-point titration of RML6 brain homogenate were adapted from (Falsig, Julius et al. 2008). ^b^ Dilutions were started from a 10 % BH. An approximate linear regression curve (y = -0.08624x + 11.63; r = 0.9388) was obtained from the mean incubation periods of dilutions 10^-1^ to 10^-7^ (Prusiner, Cochran et al. 1982, Brandner, Isenmann et al. 1996) and used to estimate the titers of the beads. Red.: titer log reduction.

| End-point titration of RML6 <sup>a</sup> |  |  | Steel bead *in vivo* assay |  |  | Decontamination |  |  |
| --- | --- | --- | --- | --- | --- | --- | --- | --- |
| RML6 <sup>b</sup><br>dilution | Mean<br>incubation<br>period<br>(days ± SD) | Titer<br>(Log LD<sub>50</sub><br>units ml<sup>-1</sup>) | RML6<br>dilution<br>to coat the<br>beads | Mean<br>incubation<br>period<br>(days ± SD) | Estimated<br>titer<br>(Log LD<sub>50</sub><br>units ml<sup>-1</sup>) | Chemical | Mean<br>incubation<br>period<br>(days ± SD) | Red.<br>(log<sub>10</sub>) |
| 10<sup>-1</sup> | 54 ± 1 | 7.9 |  |  |  | NaOH | >250 | ≥5.1 |
| 10<sup>-2</sup> | 59 ± 2 | 6.9 | 10<sup>-2</sup> | 75.9 ± 1.8 | 5.1 | 36 BS | >250 | ≥5.1 |
| 10<sup>-3</sup> | 69 ± 1 | 5.9 | 10<sup>-3</sup> | 92.8 ± 5.5 | 3.6 | 28 AO | >250 | ≥5.1 |
| 10<sup>-4</sup> | 77 ± 2 | 4.9 | 10<sup>-4</sup> | 121 ± 13.2 | 1.2 | 28 AO/TZ | >250 | ≥5.1 |
| 10<sup>-5</sup> | 86 ± 2 | 3.9 | 10<sup>-5</sup> | >250 | - | 70B | >250 | ≥5.1 |
| 10<sup>-6</sup> | 93 ± 2 | 2.9 | 10<sup>-6</sup> | >250 | - |  |  |  |
| 10<sup>-7</sup> | 119 ± 4 | 1.9 | 10<sup>-7</sup> | >250 | - |  |  |  |
| 10<sup>-8</sup> | 121, >236 | 0.9 |  |  |  |  |  |  |
| 10<sup>-9</sup> | >236 | - |  |  |  |  |  |  |

We next explored the efficiency of 70B, 28AO, 28AO/TZ, and 36BS to reduce prion infectivity on the prion coated beads in the bioassay. 30 µL of prion exposed beads (1% (w/v) RML6) treated with the different formulations or NaOH (same samples/conditions as used in the TESSA (Fig. 2A and B and Fig. S1)) were inoculated i.c. into *tg*a*20* mice in three groups of three animals. None of the inoculated mice (n= 9 per formulation) developed detectable prion disease until study termination at 250 dpi (Table 3; Fig. 3B and C). To further investigate the absence of any seeding active prions in the brains of these mice, BH samples of three individual mice per group were analyzed by a standard SAA (Wilham, Orrú et al. 2010, Frontzek, Pfammatter et al. 2016). No signal was observed for any of the brain homogenates of mice previously inoculated with treated beads, indicative for the absence of seeding active prions (Fig. S2), whereas samples from mice inoculated with untreated prion-exposed beads showed a positive signal. Our data thus indicate that all formulations and NaOH reduced prion infectivity on the prion-exposed beads below detectability in the mouse bioassay. This corresponds to a titer reduction of ≥ 5.1 log10 units ml^-1^ based on the established reference curve for prion-exposed beads. In conclusion, our data show that all tested formulations are highly effective for the inactivation of prions on medical steel surfaces.

## Discussion

In this study, we have developed TESSA, a stainless-steel bead SAA assay, as a tool for the rapid screening of a large number of chemical formulations to assess their effectiveness to inactivate prions on steel surfaces. For its establishment, we opted for the use of micron scale stainless steel beads as prion carriers to mimic the surface of surgical steel instruments, since they are more practical to handle than most other previously utilized prion carriers due to their ease of pipetting (Edgeworth, Farmer et al. 2011, Belondrade, Nicot et al. 2016, Hughson, Race et al. 2016, Mori, Atarashi et al. 2016, Williams, Hughson et al. 2019). We further air-dried the beads after prion exposure to perform the screen under most stringent conditions, since air-dried prions bind more tightly to surfaces and show a higher resistance to decontamination treatments (Secker, Hervé et al. 2011). Using this approach, our results showed that the steel beads were able to efficiently bind prions and amplify prions in a way identical to that previously described for prions coated to other carriers (Belondrade, Nicot et al. 2016, Hughson, Race et al. 2016, Mori, Atarashi et al. 2016, Williams, Hughson et al. 2019, Eraña, Pérez-Castro et al. 2020).

For the screen, we focused on investigating strong to mild alkaline (> pH 9.5) and oxidative formulations which we grouped in four classes based on their main anti-prion active ingredients. We identified 70B, a hypochlorite-based formulation, to be the most effective formulation, whereas all alkaline formulations tested in the screen showed only a moderate to no prion inactivation efficiency under conditions simulating automated washer-disinfector cleaning processes. Formulation 70B proved to be effective at a concentration of 1 % (v/v, 55 °C, 10 min) and at even lower concentrations and shorter contact times (0.5% v/v, 5 min). Additionally, we found that the alkaline formulations, 28AO (Hirata, Ito et al. 2010), 28AO/TZ, and 36BS used for the applicability testing of TESSA also fully eliminated the propagation activity of prions bound to the metal surfaces. After having identified 70B, 28AO, 28AO/TZ and 36BS as being highly efficient in TESSA, we further confirmed their effectiveness by PK immunoblotting where all formulations eliminated the amount of bead bound PrP^Sc^ below detectability.

Although *in vitro* SAAs are more widely applied for the evaluation of prion decontaminants (Pritzkow, Wagenführ et al. 2011, Belondrade, Nicot et al. 2016, Hughson, Race et al. 2016, Mori, Atarashi et al. 2016, Williams, Hughson et al. 2019, Eraña, Pérez-Castro et al. 2020), bioassays are still required to claim a new formulation as prionicidal (Taylor 2000, Fichet, Comoy et al. 2007, Giles, Glidden et al. 2008). In this study, all four formulations, along with the reference NaOH, reduced prion titers in the mouse bioassay below the detection limit of the maximum achievable sensitivity of 5.1 LD50 ml^-1^ which was determined in the reference bioassay. These results demonstrate that all formulations fulfilled or even exceeded the criteria of a log reduction of 4.5 in infectivity, required by the WHO for a prion decontaminant to be considered as effective (WHO 2000). The data for 28AO also confirm previous *in vitro* experiments (Hirata, Ito et al. 2010), and thus provide further evidence for its prionicidal efficacy. In line with previous studies (Wilham, Orrú et al. 2010, Vascellari, Orrù et al. 2012, Elder, Henderson et al. 2013, Henderson, Manca et al. 2013, Henderson, Davenport et al. 2015, Hughson, Race et al. 2016, Bélondrade, Jas-Duval et al. 2020), our results further show that the data obtained by TESSA and the mouse bioassay are well correlated, suggesting TESSA as a viable method for the screening of new prion decontaminants on prion contaminated steel surfaces. However, whether SAAs can eventually be used as a complete substitute for the biological infectivity determined by bioassays remains a critical question from a medical safety perspective and more data need to be acquired to answer this question definitively.

An important prerequisite for effective prion decontaminants is not only that they efficiently eliminate any prion contamination, but also that they are compatible with corrosion-sensitive materials to avoid damage to delicate surfaces of medical devices. Formulations 28AO, 28AO/TZ, and 36BS are gentle alkaline cleaners which are applicable to corrosion-sensitive surfaces of medical devices. This property, along with our finding for effective prion inactivation, suggests their potential application as an additional safety measure in the multistep reprocessing procedures of reusable medical devices applied in medical settings as advised by the WHO (WHO 2000) to increase the decontamination efficiency and reduce the risk of iCJD transmission through unintentionally contaminated surgical instruments.

Chlorine-based disinfectants, such as sodium hypochlorite, belong to the most effective prion decontaminants (Taylor 1993, WHO 2000, Williams, Hughson et al. 2019), but have limited relevance for routine instrument reprocessing due to their strong corrosive properties. In our work, formulation 70B, designed to have a low active chlorine concentration to meet the requirement for low corrosion properties, turned out to be a highly effective decontaminant that reduced the prion infectivity titer below detectability. However, chlorine-based products are constrained by their limited shelf life due to degradation of the active chlorine content over time (Clarkson, Moule et al. 2001, Nicoletti, Siqueira et al. 2009, Hughson, Race et al. 2016). The simultaneous decline in the active chlorine content and decontamination proficiency was also observable in our findings with formulation 70B and we therefore would advise for using either freshly prepared solutions or the addition of chlorine stabilizers. Under these conditions, formulation 70B could be a useful alternative cleaner for the decontamination of prion contaminated surfaces.

The effectiveness of prion decontaminants can also depend on the prion strain type (McDonnell and Burke 2003, Giles, Glidden et al. 2008, Edgeworth, Jackson et al. 2009, Edgeworth, Farmer et al. 2011, Edgeworth, Sicilia et al. 2011, McDonnell, Dehen et al. 2013, Ellett, Revill et al. 2020). Due the scarcity of suitable human prion mouse models (Telling, Scott et al. 1995, Watts and Prusiner 2014), we relied on the rodent-adapted prion strain, RML6, which has been widely used as a model strain for the development and validation of prion decontaminants (Fichet, Comoy et al. 2004, Lemmer, Mielke et al. 2008, Edgeworth, Sicilia et al. 2011, McDonnell, Dehen et al. 2013). However, the efficacy of prion inactivation procedures tested on rodent prions cannot be completely generalized to human prions (Adjou, Demaimay et al. 1996, Kawasaki, Kawagoe et al. 2007, Bélondrade, Jas-Duval et al. 2020). In the future, however, SAAs such as TESSA, that are applicable to many different prion strains (Hughson, Race et al. 2016, Cooper, Hoover et al. 2019, Avar, Heinzer et al. 2022), could be used as animal-free methods for the additional validation of prion decontaminants on multiple prion strains.

In conclusion, TESSA allows the detection of steel-bead adsorbed prions and represents an effective method for the rapid screening and evaluation of the effectiveness of new prion decontaminants on steel surfaces. Our data further show that formulations 28AO, 28AO/TZ, 36BS and 70B efficiently inactivate prions below the detection limit as demonstrated by several orthogonal *in vitro* methods as well as by a mouse bioassay. These formulations could therefore be suitable as mild yet effective, anti-prion decontaminants for routine instrument reprocessing to increase the safety of reusable surgical and medical instruments.

## Materials and Methods

### Chemical compositions of the formulations

Products and conditions used in this study were provided by Borer Chemie AG (Zuchwil, Switzerland) and are summarized in Tables 1 and 2. Four different main chemical categories were tested, including series of mild alkaline alkanolamine (Category 1), alkaline enzymatic phosphate/silicate (Dec TWIN PH10; Category 2), strong alkaline hydroxide-and silicate-based (deconex^®^ OP 152, deconex^®^ HT and deconex^®^ MT; Category 3), and hypochlorite-based formulations and solutions (Category 4). To improve the anti-prion and cleaning properties of the formulations, they were supplemented with different ingredients. Enzymatic detergents, such as amylases, cellulases and lipases, were added for the digestion and removal of lipids, carbohydrates, polysaccharides and fatty deposits (Olsen and Falholt 1998), whereas proteases were used to hydrolyze and remove proteins and to directly act on prions (D’Castro, Wenborn et al. 2010). In addition, non-ionic surfactants, chelating agents and surface active reagents (e.g. fatty alcohol alcoxylates and alkyl glucosides) were supplemented to induce a low-foam profile that is required for automated applications, and/or as corrosion inhibitors or wetting reagents to reduce the surface tension (Geetha and Tyagi 2012).

### Coating of the stainless-steel beads with prions and decontamination procedure

Unless otherwise specified, 100 µL of 1 % RML6 brain homogenate (in-house produced, 9.9 ± 0.2 log LD50 units g^-1^ (Falsig, Julius et al. 2008)) or NBH were incubated with 25 mg of autoclaved stainless steel beads (AISI 316L – grade surgical steel, average diameter of <20 µm; Thyssenkrupp; material number 1.4404) for 2 h at 37 °C under agitation at 1100 rpm. After incubation, the supernatant was discarded, and beads were air-dried at 37 °C for 1 h. Beads were washed four-times with sterile phosphate-buffered saline (PBS) to remove any residual RML6 using a DynaMag-2 (Thermo Fisher) for magnetic separation of the beads from the buffer and stored at −20 °C until further use. For prion-inactivation, RML6 coated beads were treated with the different formulations as specified in Tables 1 and 2. After decontamination, beads were washed 10 times with ddH2O and stored in PBS at −20 °C until further usage.

### TESSA

For the TESSA reactions, the same experimental conditions were used as previously described for a standard SAA (Atarashi, Sano et al. 2011, Frontzek, Pfammatter et al. 2016, Sorce, Nuvolone et al. 2020). Briefly, the reaction buffer of TESSA was composed of recombinant hamster PrP(23-231) (produced in-house; filtered using 100 kDa centrifugal filters (Pall Nanosep OD100C34)) at a final concentration of 0.1 mg/mL, 1 mM EDTA, 10 µM Thioflavin T, 170 mM NaCl and 1 × PBS (incl. 130 mM NaCl) (Frontzek, Pfammatter et al. 2016). To the 70 µL reaction buffer per well (96-well plate), RML6 coated beads either untreated or treated with the different formulations as described above were added (30 µL of 25 mg/mL ∼ 750 µg). The TESSA reactions were performed in a FLUOstar Omega plate reader (BMG Labtech) with cyclic shaking modes of 7 × (90 s shaking (900 rpm (double orbital), 30 s rest)), 60 s reading) at 42 °C. Reading was carried out with excitation at 450 nm and emission at 480 nm every 15 min for 105 h and reactions were performed in quadruplicates. The microplate for each run contained four control wells of each RML6 treated with ddH2O, and of RML6 coated beads treated with 1 M NaOH. Only fluorescence positive reactions between 0 and 83 h hours were considered for the data evaluation, because of an increasing occurrence of spontaneous aggregation events at later reaction times.

For the preparation of the *tga*20 brain homogenate of the mice inoculated with the beads, brains were homogenized in 0.32 M sucrose (10 % (w/v), Sigma) using the Precellys 24 (Bertin Instruments) as previously described (Frontzek, Pfammatter et al. 2016) and diluted 20’000 times in PBS. A total of 98 µL of reaction buffer was dispensed into a well of a 96-well plate and 2 µL of each BH sample was added. The assay was then performed under the conditions as described above.

### Immunoblotting

Samples containing beads were prepared as mentioned above. As controls, NBH or RML6 brain homogenates were used after determination of total protein levels with a bicinchoninic acid assay (BCA, Pierce) according to the manufacturer and were adjusted to 20 µg of protein and PK digested. To assess PK-resistant PrP^Sc^ levels on beads, 20 µL PBS with PK (25 µg/mL) was added to the beads and the PK digestion was performed directly on the beads at 37 °C for 30 minutes under shaking at 1’100 rpm. To stop the digestion, 4 × LDS containing loading dye (NuPAGE, Thermo Fisher) and 1 mM 1,4-dithiothreitol (Roche) were added prior to boiling the samples at 95 °C for 10 minutes and centrifugation at 2’000 rpm for 5 minutes. Samples were loaded on a 12 % Bis-Tris Gel (NuPAGE, Thermo Fisher) and transferred to a nitrocellulose membrane using the iBlot system (Thermo Fisher). Membranes were blocked with 5 % SureBlock (LuBio science) and probed with 1:10’000 monoclonal POM1 antibody (Polymenidou, Moos et al. 2008) in 1 % Sure-Block/PBS with 0.1 % Tween-20 (PBS-T). As a detection antibody, an HRP-conjugated goat anti-mouse antibody (1:10’000, Invitrogen) was used in 1 % SureBlock-PBS-T. Membranes were developed with Crescendo HRP substrate (Millipore) and imaging was done using the LAC3000 system (Fuji).

### Mouse bioassay in *tg*a*20* mice

All animal experimentation was performed under compliance with the rules and regulations by the Swiss Confederation on the Protection of Animal Rights. All protocols used in this study were approved by the Animal Welfare Committee of the Canton of Zurich under the permit number ZH040/15. *Tg*a*20* mice were kept under general anesthesia after isoflurane treatment and inoculated i.c. using 30 µL of a suspension of either NBH or RML6 coated beads that were either untreated or treated with the different decontaminants as described above. Inoculated mice were subjected to health monitoring every second day until the appearance of terminal clinical symptoms of scrapie. On the first day the animals presented themselves as being terminally sick, they were sacrificed under isoflurane anesthesia. Brains of mice were dissected and analyzed immunohistochemically for the presence of a prion disease. Histological analysis was performed on one hemisphere of the dissected brains after inactivation in 96 % formic acid and fixation in formalin. Hematoxylin and eosin stain (H&E) was used to confirm spongiform changes in brain tissue sections, whereas SAF84 staining was used to visualize PrP^Sc^ aggregates.

## Acknowledgements

We thank Linda Irpinio and Dezirae Schneider for technical assistance. This work was funded by Innosuisse - the Swiss Innovation Agency.

## Author Contributions

**Conceptualization:** UR AA SH.

**Formal analysis:** DH MA RM PS.

**Funding acquisition:** UR AA SH.

**Investigation:** DH MA RM PS.

**Project administration:** AA SH.

**Resources:** MTB BK SM UR AA SH.

**Supervision:** AA SH.

**Visualization:** DH MA MTB BK SM SH.

**Writing – original draft:** DH MA SH.

**Writing – review & editing:** all authors

## Supplemental Information Supplemental Figures

**Supplemental Figure 1:**
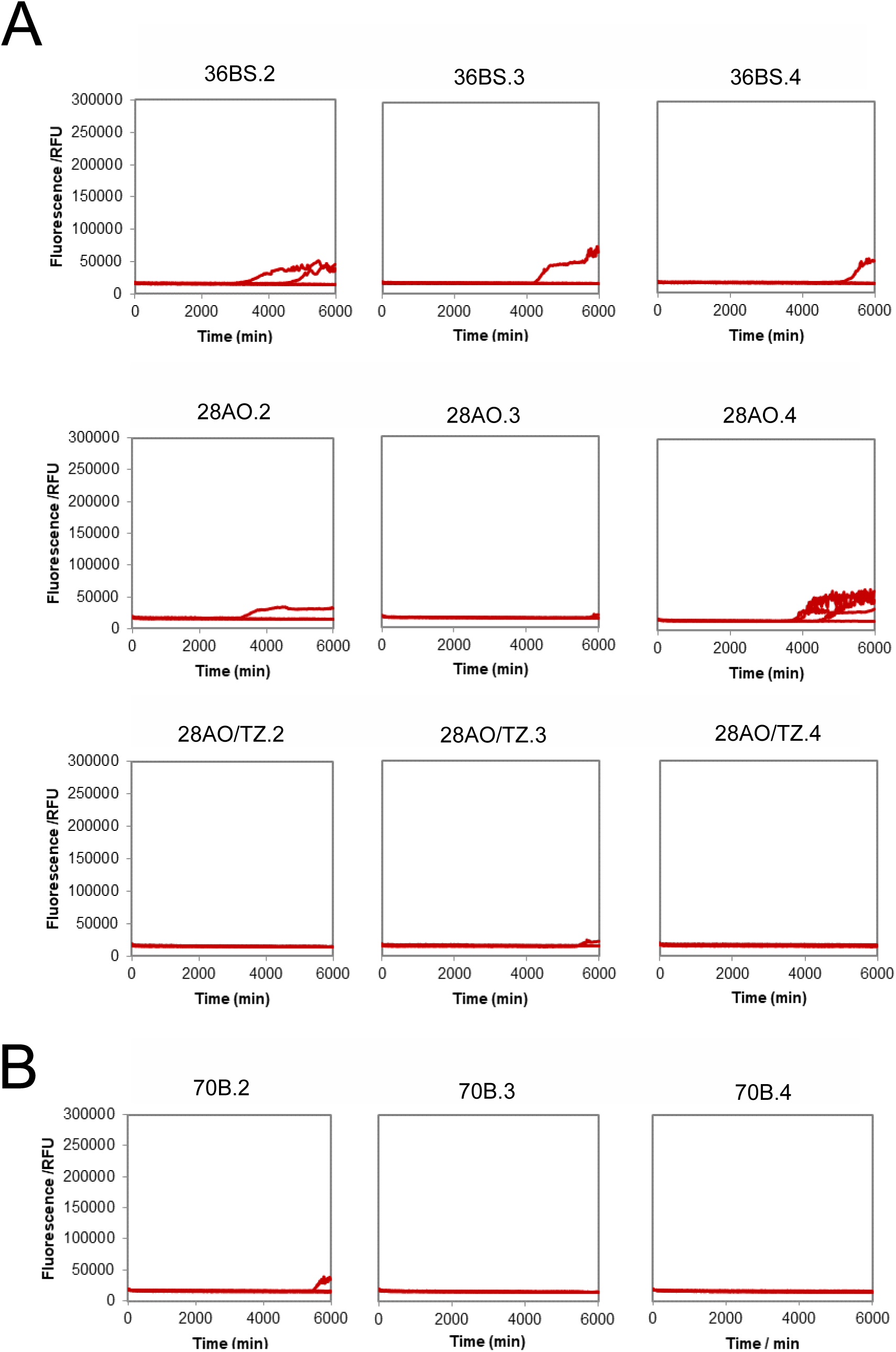
Additional TESSA replicates for the confirmation of the decontamination efficacy of the four formulations. **(A)** Same as shown in Figure 2A, but for three further TESSA replicates of RML6 exposed beads after decontamination with formulation 28AO, 28AO/TZ and 36BS. Shown are three reactions for each condition in quadruplicates. (B) Same as shown in Figure 2B, but for three further replicates of RML6 exposed beads after decontamination with formulation 70B. Shown are three reactions for each condition in quadruplicates.

**Supplemental Figure 2:**
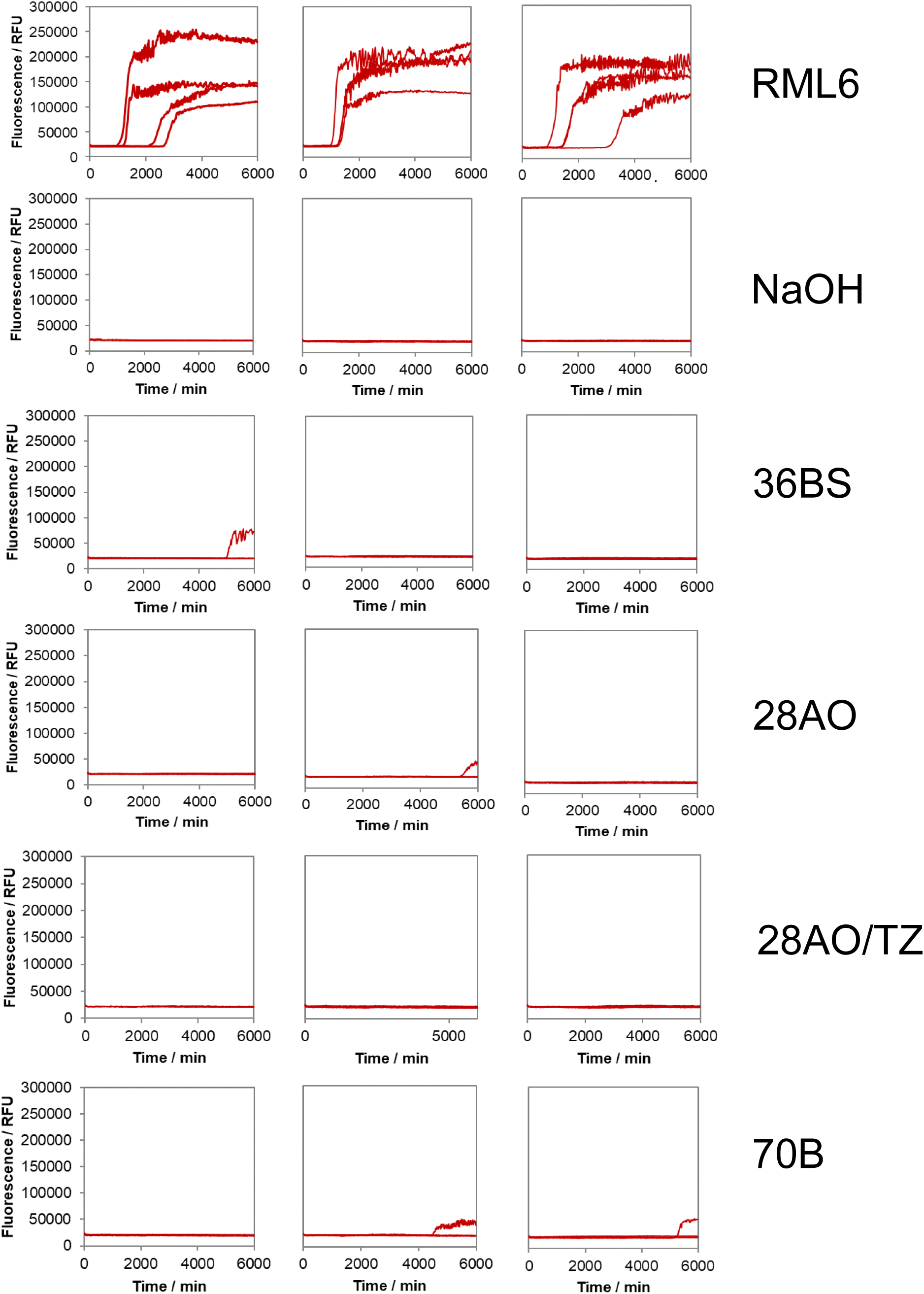
Standard SAA analysis of BHs from *tg*a*20* mice inoculated with prion-contaminated steel beads treated with the different decontaminants. SAA analysis of BHs of three individual mice per condition. *Tg*a*20* mice were inoculated with steel-beads previously exposed to RML6 and treated with ddH2O (RML6), 1M NaOH or different formulations and brains were assessed for their seeding activity.

**Source data.**
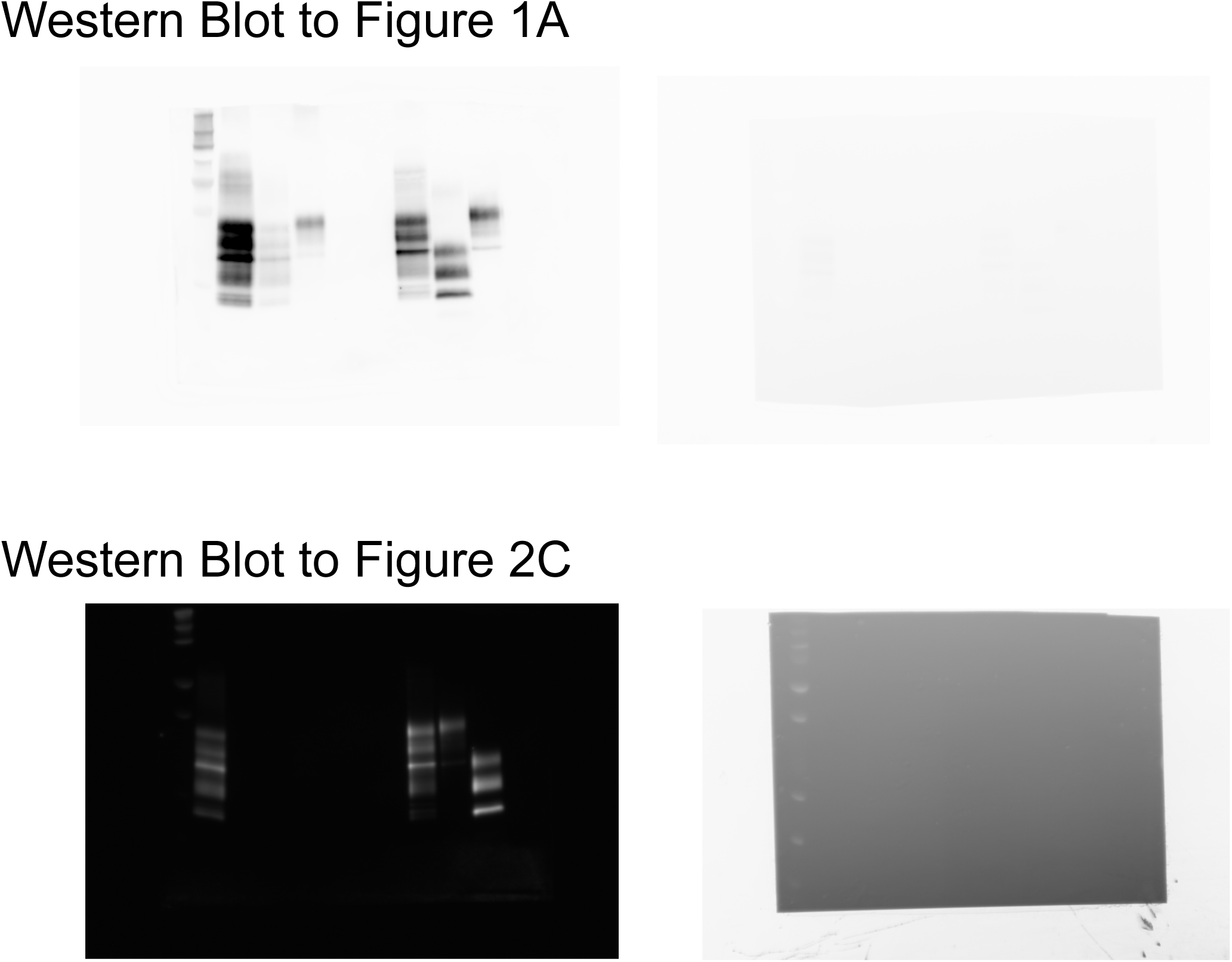
**(A)** Source data of Figure 1A. Left panel: POM1 immunoblot, right panel: marker. **(B)** Source data of Figure 2C. Left panel: POM1 immunoblot, right panel: marker.

## References

1. Adjou, K. T., R. Demaimay, C. I. Lasmézas, M. Seman, J. P. Deslys and D. Dormont (1996). “Differential effects of a new amphotericin B derivative, MS-8209, on mouse BSE and scrapie: implications for the mechanism of action of polyene antibiotics.” Res Virol 147(4): 213–218.

2. Aguzzi, A. and A. M. Calella (2009). “Prions: protein aggregation and infectious diseases.” Physiol Rev 89(4): 1105–1152.

3. Aguzzi, A., M. Heikenwalder and M. Polymenidou (2007). “Insights into prion strains and neurotoxicity.” Nat Rev Mol Cell Biol 8(7): 552–561.

4. Alper, T., W. A. Cramp, D. A. Haig and M. C. Clarke (1967). “Does the agent of scrapie replicate without nucleic acid?” Nature 214(5090): 764–766.

5. Atarashi, R., K. Sano, K. Satoh and N. Nishida (2011). “Real-time quaking-induced conversion: a highly sensitive assay for prion detection.” Prion 5(3): 150–153.

6. Avar, M., D. Heinzer, A. M. Thackray, Y. Liu, M. Hruska-Plochan, S. Sellitto, E. Schaper, D. P. Pease, J. A. Yin, A. K. Lakkaraju, M. Emmenegger, M. Losa, A. Chincisan, S. Hornemann, M. Polymenidou, R. Bujdoso and A. Aguzzi (2022). “An arrayed genome-wide perturbation screen identifies the ribonucleoprotein Hnrnpk as rate-limiting for prion propagation.” EMBO J: e112338.

7. Bélondrade, M., C. Jas-Duval, S. Nicot, L. Bruyère-Ostells, C. Mayran, L. Herzog, F. Reine, J. M. Torres, C. Fournier-Wirth, V. Béringue, S. Lehmann and D. Bougard (2020). “Correlation between Bioassay and Protein Misfolding Cyclic Amplification for Variant Creutzfeldt-Jakob Disease Decontamination Studies.” mSphere 5(1).

8. Belondrade, M., S. Nicot, V. Béringue, J. Coste, S. Lehmann and D. Bougard (2016). “Rapid and Highly Sensitive Detection of Variant Creutzfeldt-Jakob Disease Abnormal Prion Protein on Steel Surfaces by Protein Misfolding Cyclic Amplification: Application to Prion Decontamination Studies.” PLoS One 11(1): e0146833.

9. Berberidou, C., K. Xanthopoulos, I. Paspaltsis, A. Lourbopoulos, E. Polyzoidou, T. Sklaviadis and I. Poulios (2013). “Homogenous photocatalytic decontamination of prion infected stainless steel and titanium surfaces.” Prion 7(6): 488–495.

10. Bernoulli, C., J. Siegfried, G. Baumgartner, F. Regli, T. Rabinowicz, D. C. Gajdusek and C. J. Gibbs, Jr. (1977). “Danger of accidental person-to-person transmission of Creutzfeldt-Jakob disease by surgery.” Lancet 1(8009): 478–479.

11. Bolton, D. C., R. K. Meyer and S. B. Prusiner (1985). “Scrapie PrP 27-30 is a sialoglycoprotein.” J Virol 53(2): 596–606.

12. Bonda, D. J., S. Manjila, P. Mehndiratta, F. Khan, B. R. Miller, K. Onwuzulike, G. Puoti, M. L. Cohen, L. B. Schonberger and I. Cali (2016). “Human prion diseases: surgical lessons learned from iatrogenic prion transmission.” Neurosurg Focus 41(1): E10.

13. Brandner, S., S. Isenmann, A. Raeber, M. Fischer, A. Sailer, Y. Kobayashi, S. Marino, C. Weissmann and A. Aguzzi (1996). “Normal host prion protein necessary for scrapie-induced neurotoxicity.” Nature 379(6563): 339–343.

14. Caughey, B., G. J. Raymond and R. A. Bessen (1998). “Strain-dependent differences in beta-sheet conformations of abnormal prion protein.” J Biol Chem 273(48): 32230–32235.

15. Clarkson, R. M., A. J. Moule and H. M. Podlich (2001). “The shelf-life of sodium hypochlorite irrigating solutions.” Aust Dent J 46(4): 269–276.

16. Collins, S. and C. L. Masters (1996). “Iatrogenic and zoonotic Creutzfeldt-Jakob disease: the Australian perspective.” Med J Aust 164(10): 598–602.

17. Cooper, S. K., C. E. Hoover, D. M. Henderson, N. J. Haley, C. K. Mathiason and E. A. Hoover (2019). “Detection of CWD in cervids by RT-QuIC assay of third eyelids.” PLoS One 14(8): e0221654.

18. D’Castro, L., A. Wenborn, N. Gros, S. Joiner, S. Cronier, J. Collinge and J. D. Wadsworth (2010). “Isolation of proteinase K-sensitive prions using pronase E and phosphotungstic acid.” PLoS One 5(12): e15679.

19. Duffy, P., J. Wolf, G. Collins, A. G. DeVoe, B. Streeten and D. Cowen (1974). “Letter: Possible person-to-person transmission of Creutzfeldt-Jakob disease.” N Engl J Med 290(12): 692–693.

20. Edgeworth, J. A., M. Farmer, A. Sicilia, P. Tavares, J. Beck, T. Campbell, J. Lowe, S. Mead, P. Rudge, J. Collinge and G. S. Jackson (2011). “Detection of prion infection in variant Creutzfeldt-Jakob disease: a blood-based assay.” Lancet 377(9764): 487–493.

21. Edgeworth, J. A., G. S. Jackson, A. R. Clarke, C. Weissmann and J. Collinge (2009). “Highly sensitive, quantitative cell-based assay for prions adsorbed to solid surfaces.” Proc Natl Acad Sci U S A 106(9): 3479–3483.

22. Edgeworth, J. A., A. Sicilia, J. Linehan, S. Brandner, G. S. Jackson and J. Collinge (2011). “A standardized comparison of commercially available prion decontamination reagents using the Standard Steel-Binding Assay.” J Gen Virol 92(Pt 3): 718–726.

23. Elder, A. M., D. M. Henderson, A. V. Nalls, J. M. Wilham, B. W. Caughey, E. A. Hoover, A. E. Kincaid, J. C. Bartz and C. K. Mathiason (2013). “In vitro detection of prionemia in TSE-infected cervids and hamsters.” PLoS One 8(11): e80203.

24. Ellett, L. J., Z. T. Revill, Y. Q. Koo and V. A. Lawson (2020). “Strain variation in treatment and prevention of human prion diseases.” Prog Mol Biol Transl Sci 175: 121–145.

25. Eraña, H., M. Pérez-Castro, S. García-Martínez, J. M. Charco, R. López-Moreno, C. M. Díaz-Dominguez, T. Barrio, E. González-Miranda and J. Castilla (2020). “A Novel, Reliable and Highly Versatile Method to Evaluate Different Prion Decontamination Procedures.” Front Bioeng Biotechnol 8: 589182.

26. Falsig, J., C. Julius, I. Margalith, P. Schwarz, F. L. Heppner and A. Aguzzi (2008). “A versatile prion replication assay in organotypic brain slices.” Nat Neurosci 11(1): 109–117.

27. Fichet, G., E. Comoy, C. Dehen, L. Challier, K. Antloga, J. P. Deslys and G. McDonnell (2007). “Investigations of a prion infectivity assay to evaluate methods of decontamination.” J Microbiol Methods 70(3): 511–518.

28. Fischer, M., T. Rulicke, A. Raeber, A. Sailer, M. Moser, B. Oesch, S. Brandner, A. Aguzzi and C. Weissmann (1996). “Prion protein (PrP) with amino-proximal deletions restoring susceptibility of PrP knockout mice to scrapie.” EMBO J 15(6): 1255–1264.

29. Flechsig, E., I. Hegyi, M. Enari, P. Schwarz, J. Collinge and C. Weissmann (2001). “Transmission of scrapie by steel-surface-bound prions.” Mol Med 7(10): 679–684.

30. Frontzek, K., M. Pfammatter, S. Sorce, A. Senatore, P. Schwarz, R. Moos, K. Frauenknecht, S. Hornemann and A. Aguzzi (2016). “Neurotoxic Antibodies against the Prion Protein Do Not Trigger Prion Replication.” PLoS One 11(9): e0163601.

31. Geetha, D. and R. Tyagi (2012). “Alkyl Poly Glucosides (APGs) Surfactants and Their Properties: A Review.” Tenside Surfactants Detergents 49(5): 417–427.

32. Giles, K., D. V. Glidden, R. Beckwith, R. Seoanes, D. Peretz, S. J. DeArmond and S. B. Prusiner (2008). “Resistance of bovine spongiform encephalopathy (BSE) prions to inactivation.” PLoS Pathog 4(11): e1000206.

33. Head, M. W. and J. W. Ironside (2007). “vCJD and the gut: implications for endoscopy.” Gut 56(1): 9–11.

34. Henderson, D. M., K. A. Davenport, N. J. Haley, N. D. Denkers, C. K. Mathiason and E. A. Hoover (2015). “Quantitative assessment of prion infectivity in tissues and body fluids by real-time quaking-induced conversion.” J Gen Virol 96(Pt 1): 210–219.

35. Henderson, D. M., M. Manca, N. J. Haley, N. D. Denkers, A. V. Nalls, C. K. Mathiason, B. Caughey and E. A. Hoover (2013). “Rapid antemortem detection of CWD prions in deer saliva.” PLoS One 8(9): e74377.

36. Hirata, Y., H. Ito, T. Furuta, K. Ikuta and A. Sakudo (2010). “Degradation and destabilization of abnormal prion protein using alkaline detergents and proteases.” Int J Mol Med 25(2): 267–270.

37. Hughson, A. G. B. Race, A. Kraus, L. R. Sangaré, L. Robins, B. R. Groveman, E. Saijo, K. Phillips, L. Contreras, V. Dhaliwal, M. Manca, G. Zanusso, D. Terry, J. F. Williams and B. Caughey (2016). “Inactivation of Prions and Amyloid Seeds with Hypochlorous Acid.” PLoS Pathog 12(9): e1005914.

38. Kawasaki, Y., K. Kawagoe, C. J. Chen, K. Teruya, Y. Sakasegawa and K. Doh-ura (2007). “Orally administered amyloidophilic compound is effective in prolonging the incubation periods of animals cerebrally infected with prion diseases in a prion strain-dependent manner.“ J Virol 81(23): 12889–12898.

39. Lehmann, S., M. Pastore, C. Rogez-Kreuz, M. Richard, M. Belondrade, G. Rauwel, F. Durand, R. Yousfi, J. Criquelion, P. Clayette and A. Perret-Liaudet (2009). “New hospital disinfection processes for both conventional and prion infectious agents compatible with thermosensitive medical equipment.” J Hosp Infect 72(4): 342–350.

40. Lemmer, K., M. Mielke, C. Kratzel, M. Joncic, M. Oezel, G. Pauli and M. Beekes (2008). “Decontamination of surgical instruments from prions. II. In vivo findings with a model system for testing the removal of scrapie infectivity from steel surfaces.” J Gen Virol 89(Pt 1): 348–358.

41. Lipscomb, I. P., H. E. Pinchin, R. Collin, K. Harris and C. W. Keevil (2006). “Are surgical stainless steel wires used for intracranial implantation of PrPsc a good model of iatrogenic transmission from contaminated surgical stainless steel instruments after cleaning?” J Hosp Infect 64(4): 339–343.

42. Luhr, K. M., P. Low, A. Taraboulos, T. Bergman and K. Kristensson (2009). “Prion adsorption to stainless steel is promoted by nickel and molybdenum.” J Gen Virol 90(Pt 11): 2821–2828.

43. McDonnell, G. and P. Burke (2003). “The challenge of prion decontamination.” Clin Infect Dis 36(9): 1152–1154.

44. McDonnell, G., C. Dehen, A. Perrin, V. Thomas, A. Igel-Egalon, P. A. Burke, J. P. Deslys and E. Comoy (2013). “Cleaning, disinfection and sterilization of surface prion contamination.” J Hosp Infect 85(4): 268–273.

45. McKinley, M. P., D. C. Bolton and S. B. Prusiner (1983). “A protease-resistant protein is a structural component of the scrapie prion.” Cell 35(1): 57–62.

46. McKinley, M. P., R. K. Meyer, L. Kenaga, F. Rahbar, R. Cotter, A. Serban and S. B. Prusiner (1991). “Scrapie prion rod formation in vitro requires both detergent extraction and limited proteolysis.” J Virol 65(3): 1340–1351.

47. Mori, T., R. Atarashi, K. Furukawa, H. Takatsuki, K. Satoh, K. Sano, T. Nakagaki, D. Ishibashi, K. Ichimiya, M. Hamada, T. Nakayama and N. Nishida (2016). “A direct assessment of human prion adhered to steel wire using real-time quaking-induced conversion.” Sci Rep 6: 24993.

48. Nakano, Y., N. Akamatsu, T. Mori, K. Sano, K. Satoh, T. Nagayasu, Y. Miyoshi, T. Sugio, H. Sakai, E. Sakae, K. Ichimiya, M. Hamada, T. Nakayama, Y. Fujita, K. Yanagihara and N. Nishida (2016). “Sequential Washing with Electrolyzed Alkaline and Acidic Water Effectively Removes Pathogens from Metal Surfaces.” PLoS One 11(5): e0156058.

49. Nicoletti, M. A., E. L. Siqueira, A. C. Bombana and G. G. Oliveira (2009). “Shelf-life of a 2.5% sodium hypochlorite solution as determined by Arrhenius equation.” Braz Dent J 20(1): 27–31.

50. Olsen, H. S. and P. Falholt (1998). “The role of enzymes in modern detergency.” Journal of Surfactants and Detergents 1(4): 555–567.

51. Peretz, D., S. Supattapone, K. Giles, J. Vergara, Y. Freyman, P. Lessard, J. G. Safar, D. V. Glidden, C. McCulloch, H. O. Nguyen, M. Scott, S. J. Dearmond and S. B. Prusiner (2006). “Inactivation of prions by acidic sodium dodecyl sulfate.” J Virol 80(1): 322–331.

52. Polymenidou, M., R. Moos, M. Scott, C. Sigurdson, Y. Z. Shi, B. Yajima, I. Hafner-Bratkovic, R. Jerala, S. Hornemann, K. Wuthrich, A. Bellon, M. Vey, G. Garen, M. N. James, N. Kav and A. Aguzzi (2008). “The POM monoclonals: a comprehensive set of antibodies to non-overlapping prion protein epitopes.” PLoS ONE 3(12): e3872.

53. Pritzkow, S., K. Wagenführ, M. L. Daus, S. Boerner, K. Lemmer, A. Thomzig, M. Mielke and M. Beekes (2011). “Quantitative detection and biological propagation of scrapie seeding activity in vitro facilitate use of prions as model pathogens for disinfection.” PLoS One 6(5): e20384.

54. Prusiner, S. B. (1982). “Novel proteinaceous infectious particles cause scrapie.” Science 216(4542): 136–144.

55. Prusiner, S. B., S. P. Cochran, D. F. Groth, D. E. Downey, K. A. Bowman and H. M. Martinez (1982). “Measurement of the scrapie agent using an incubation time interval assay.” Ann Neurol 11(4): 353–358.

56. Saborio, G. P., B. Permanne and C. Soto (2001). “Sensitive detection of pathological prion protein by cyclic amplification of protein misfolding.” Nature 411(6839): 810–813.

57. Sandin, S., R. K. B. Karlsson and A. Cornell (2015). “Catalyzed and Uncatalyzed Decomposition of Hypochlorite in Dilute Solutions.” Industrial & Engineering Chemistry Research 54(15): 3767–3774.

58. Secker, T. J., R. Hervé and C. W. Keevil (2011). “Adsorption of prion and tissue proteins to surgical stainless steel surfaces and the efficacy of decontamination following dry and wet storage conditions.” J Hosp Infect 78(4): 251–255.

59. Sorce, S., M. Nuvolone, G. Russo, A. Chincisan, D. Heinzer, M. Avar, M. Pfammatter, P. Schwarz, M. Delic, M. Müller, S. Hornemann, D. Sanoudou, C. Scheckel and A. Aguzzi (2020). “Genome-wide transcriptomics identifies an early preclinical signature of prion infection.” PLoS Pathog 16(6): e1008653.

60. Taylor, D. M. (1993). “Inactivation of SE agents.” Br Med Bull 49(4): 810–821.

61. Taylor, D. M. (2000). “Inactivation of transmissible degenerative encephalopathy agents: A review.” Vet J 159(1): 10–17.

62. Telling, G. C., M. Scott, J. Mastrianni, R. Gabizon, M. Torchia, F. E. Cohen, S. J. DeArmond and S. B. Prusiner (1995). “Prion propagation in mice expressing human and chimeric PrP transgenes implicates the interaction of cellular PrP with another protein.” Cell 83(1): 79–90.

63. Thomas, J. G., C. E. Chenoweth and S. E. Sullivan (2013). “Iatrogenic Creutzfeldt-Jakob disease via surgical instruments.” J Clin Neurosci 20(9): 1207–1212.

64. Vascellari, S., C. D. Orrù, A. G. Hughson, D. King, R. Barron, J. M. Wilham, G. S. Baron, B. Race, A. Pani and B. Caughey (2012). “Prion seeding activities of mouse scrapie strains with divergent PrPSc protease sensitivities and amyloid plaque content using RT-QuIC and eQuIC.” PLoS One 7(11): e48969.

65. Watts, J. C. and S. B. Prusiner (2014). “Mouse models for studying the formation and propagation of prions.” J Biol Chem 289(29): 19841–19849.

66. WHO (2000). WHO Infection Control Guidelines for Transmissible Spongiform Encephalopathies. WHO.

67. Wilham, J. M., C. D. Orrú, R. A. Bessen, R. Atarashi, K. Sano, B. Race, K. D. Meade-White, L. M. Taubner, A. Timmes and B. Caughey (2010). “Rapid end-point quantitation of prion seeding activity with sensitivity comparable to bioassays.” PLoS Pathog 6(12): e1001217.

68. Williams, K., A. G. Hughson, B. Chesebro and B. Race (2019). “Inactivation of chronic wasting disease prions using sodium hypochlorite.” PLoS One 14(10): e0223659.

69. Zobeley, E., E. Flechsig, A. Cozzio, M. Enari and C. Weissmann (1999). “Infectivity of scrapie prions bound to a stainless steel surface.” Mol Med 5(4): 240–243.

